# Metabolic diversity of bacteria and yeast from commercial probiotic products illustrated by phenotypic profiling

**DOI:** 10.1101/364687

**Authors:** Juliana M. Ansari, Christine Colasacco, Elli Emmanouil, Scott Kohlhepp, Olivia Harriott

## Abstract

The number of probiotic products in the marketplace is on the rise, gaining momentum along with the upsurge in research on the role of the human gut microbiome in health. Although such products are considered safe for consumption, these probiotic supplements and beverages are not subject to stringent federal regulation for quality. While only certain strains of probiotic microbes have been studied for efficacy in clinical trials, the ingredient labels of commercial probiotics do not always list the strain names. In this study, we investigated the diversity of the bacteria and yeast sold in these products. From a representative selection of commercially available probiotic supplements and beverages, we cultured microbes and identified them with standard methods (16S rRNA gene sequencing, mass spectrometric identification, and Biolog phenotypic profiling), then assessed whether there were strain-specific differences in nutrient metabolism and tolerance to compounds across the isolates from different products. *Bacillus coagulans, Bacillus subtilis, Lactobacillus plantarum, Lactobacillus rhamnosus,* and the yeast *Saccharomyces boulardii* were cultured from 21 commercial probiotic products (fifteen probiotic supplements and six probiotic beverages). Phenotypic profiling revealed metabolic diversity in carbon source usage and tolerance to compounds, within species from different probiotics and from environmental isolates of strains belonging to the same species. Despite this strain level diversity, we observed that up to half of the probiotic supplements for sale in retail and drugstores only list the species, but not the specific strain, on the label. This study highlights that existing labeling conventions for probiotics are insufficient to convey the strain identity and diversity in these products, underscoring the need for clear strain identification and verification of strain-specific probiotic properties, particularly when moving toward therapeutic applications of beneficial microbes.

## Introduction

A widespread awakening in the public and medical community’s interest in commensal bacteria for promoting health is currently underway. Accelerated by the ease and affordability of rapid DNA sequencing technology, an avalanche of studies in animal models and humans has linked the microorganisms living on or in the body to countless health conditions. In the decade since the Human Microbiome Project was initiated to understand how microbes impact human physiology and disease [1], hundreds of studies are published each year exploring the roles of the microbiome (the microbial community inhabiting the human body) in a wide range of diseases. The combination of impacts on inflammation, immunity, nutrient metabolism, and even behavior may inextricably tie the healthy balance vs. imbalance (dysbiosis) of the human gut microbiome to one’s overall functioning as a healthy human organism. While the composition of bacterial (and fungal, viral, and protozoan) residents of the body is being mapped with increasing phylogenetic detail, many of the organisms are novel and the biological functions of many of their genes remain unknown [2].

This tidal wave of studies on the human microbiome has kindled renewed interest in the idea of adding beneficial bacteria (probiotics) or foods that selectively enhance growth of certain beneficial microbes (prebiotics) to the diet. Supplementing food with microorganisms is by no means new, with fermented foods and beverages common in the diets of cultures around the world, yet mainstream interest and market demand for probiotic foods is growing in parallel with research on the microbiome, nutrition, and health [3]. The promoted advantage of probiotics is the maintenance or restoration of the balance between pathogens and healthy necessary bacteria, via mechanisms such as binding to pathogens, competition for nutrients, antimicrobial production, and modulating the immune system [4]. In addition to their established roles in female reproductive health [5], probiotics have been recognized for myriad effects on digestive health [6].

Obesity has been shown to be correlated with the microbiota, linked to an imbalance in energy homeostasis, and probiotics and/or prebiotics show potential to address this [7]. According to Salazar et al., prebiotic ingestion by obese women caused an increase in *Bifidobacterium* species, attenuated short-chain fatty acid (SCFA) production and thus abated metabolic factors correlated with obesity [8]. Additionally, the consumption of probiotics has been demonstrated to improve insulin resistance syndrome, type 2-diabetes and non-alcoholic fatty liver disease [6]. **ADD_Science paper T2D.** There is evidence that probiotics assist with the production of vitamin B and necessary organic acids and amino acids, with host absorption of vitamins and minerals, and with the production of enzymes such as esterase, lipase and co-enzymes essential to metabolic processes [6]. The composition of the microbiota is also associated with irritable bowel syndrome (IBS), a chronic disorder associated with abdominal pain, distention and abnormal bowel movements, along with low grade inflammation and alterations to the gut immune system [9]. Substantial evidence indicates the efficacy of specific probiotics for alleviating the symptoms of IBS [10]. Other gastrointestinal disorders such as traveler’s diarrhea [11] and antibiotic-associated diarrhea [12], have been shown to be treated or prevented with the introduction of probiotics.

Still, the health claims made by many probiotic foods and supplements are often many steps ahead of the science backing the studies. Despite a growing body of clinical trials supporting the specific benefits of well-established strains [10], more carefully designed and controlled studies are needed [13][14]. This gap between the advertised benefits of probiotics and the evidence to support their efficacy, is due in part to the limited regulation on probiotic supplements, which are categorized by the Food and Drug Administration (FDA) as food additives or ingredients rather than drugs. The microbial strains they contain are classified as “GRAS” (generally recognized as safe), but the health claims on packages are not verified by the FDA. With research on beneficial bacteria advancing rapidly, it is expected that if the promise of microbiome-based therapies lives up to projections, then ‘precision probiotics’ aiming to prevent or treat certain health conditions may soon be classified as drugs and subjected to more stringent regulatory scrutiny and burden of proof for efficacy. Indeed, the end of the 2010s may represent the calm before the storm of ‘next-generation probiotics’ or ‘bugs as drugs,’ discovered from research on healthy microbiomes, that could be used to treat specific diseases.

When probiotics are evaluated for their ability to treat specific conditions in clinical trials, the strains used have demonstrated probiotic properties, yet the strain-level identity of probiotic bacteria is not always provided on the ingredient label [15]. While in some cases probiotic properties are species- or genus-wide [16], this omission raises several concerns, with safety being the foremost. Secondly, for those brands that do not list the strain, it is possible that the included strain does not actually possess the probiotic effects of clinically verified strains. Variability in cultivation and processing during probiotic manufacturing, also raises the possibility that the integrity of the particular probiotic strain is lost over time, through continuous passaging in industrial fermentations. When bacterial populations are cultured repeatedly over time, new stable genetic populations can emerge, such as in ongoing experimental evolution studies of *Escherichia coli* [17] and *Burkholderia cenocepacia* following repeated transfers of the same starting population [18]. It is possible that this could occur after repeated passaging of fast-growing industrial microorganisms, perhaps selecting for better growth under industrial fermentation conditions, and potentially leading to a reduction in their probiotic properties.

Identifying microbes at the strain level and ensuring that their beneficial properties are not lost during the culturing and manufacturing process, is imperative for the live organisms to exert their reported effect. The current study aimed to test how well routine genetic and phenotypic methods of microbial identification could distinguish the microbes in common commercial probiotics at the species and strain level, and to investigate the metabolic differences between the bacterial strains in each product. Three approaches were used to identify each probiotic isolate: amplification and sequencing of the 16S rRNA gene, Matrix Assisted Laser Desorption Ionization Time-of-Flight (MALDI-TOF) mass spectrometry, and the Biolog Microbial Identification system, a 96-well plate multiplex phenotypic assay consisting of 71 carbon source utilization tests and 23 chemical sensitivity tests [19]. We hypothesized that comparing the phenotypic profiles of these common probiotics would reveal differences in their nutrient metabolism (both across different probiotic species, and among strains of the same species). We also explored the prevalence of certain types of bacteria in the marketplace and compared this to the quantity of published research supporting their beneficial effects.

## Methods

### Isolation of bacteria from commercial probiotic products

Probiotic products were purchased in 2016-2017, opened within one month of the date of purchase and bottles were stored at 4 °C. “Single-strain” probiotics contained only one species of live microorganism, and “Multi-strain” probiotics contained two or more species listed on the label; however only one isolated microbe was identified from each product. For those in capsule form, the capsule was aseptically emptied into a microcentrifuge tube containing 1 ml of sterile water and mixed thoroughly. Tablets were ground with a sterilized mortar and pestle and combined with sterile water. Probiotic beverages were sampled using sterile swabs, directly from the original bottle. Using a sterile inoculating loop, the sample was streaked for isolation onto the surface of the appropriate agar growth medium. For culturing *Lactobacillus* species and *Bacillus coagulans,* MRS (de Man, Rogosa, and Sharpe) agar was used, and TSA (Tryptic Soy Agar) was used for *Bacillus subtilis*. Yeast were cultured on SDA (Sabouraud Dextrose Agar) or MRS. Agar plates were incubated at 30-33 °C for 48-72 hours aerobically, and individual isolated colonies were identified. For single strain products, a well-isolated colony was inoculated into liquid medium and incubated for 24 hours, then used to make a frozen glycerol stock by mixing 700 μl of the overnight culture with 300 μl of sterile 50% glycerol (final concentration 15% glycerol) and stored at −80 °C. For multi-strain products, one colony type was selected, and re-streaked for isolation prior to storing as a frozen glycerol stock.

To observe cells from isolated colonies, isolated bacteria were Gram stained following the standard procedure [20] and viewed under oil immersion with the 100x objective lens and photographed. As a preliminary differentiation step between *Lactobacillus* and *Bacillus,* endospore staining was performed on 1-week old cultures following the Schaeffer-Fulton procedure (Harley, 2017). Heat fixed smears were covered with a small piece of paper towel, and placed on a rack over a steaming water bath. Malachite green dye (5%) was added on top of paper and heated for 5-7 minutes, adding fresh stain as liquid evaporated. Slides were rinsed in water, counterstained with safranin for 1 minute, then viewed under oil immersion with the 100x objective lens and photographed.

### Antibiotic susceptibility testing

The Kirby-Bauer test for antibiotic susceptibility was followed with minor modifications. The typical medium for this assay is Mueller Hinton, however Mueller Hinton was used for *Bacillus subtilis* and MRS agar used for *Lactobacillus* spp. Using sterile swabs, overnight liquid cultures were spread in a zig-zag pattern to create a “lawn” of growth on large 150-mm agar plates. A 12-place BD BBL Sensi-Disc Dispenser was used to deposit the following antibiotic susceptibility Sensi-discs (BD, Franklin Lakes, NJ) onto the agar surface: Ampicillin (AM10), Bacitracin (B10), Chloramphenicol (C30), Ciprofloxacin (CIP5), Erythromycin (E15), Gentamicin (GM10), Kanamycin (K30), Neomycin (N30), Penicillin (P10), Streptomycin (S10), Tetracycline (Te30), and Vancomycin (VA30). After 24 hours of incubation at 33 °C, zones of inhibition were measured and compared to the known diameter sizes for susceptibility on the reference Zone Diameter Interpretive Chart, updated by the National Committee for Clinical Laboratory Standards, accessed in (Harley, 2017). Susceptibility to each antibiotic was recorded as susceptible (S), resistant (R), or intermediate (I) based on the diameter of the zone of inhibition.

### BIOLOG Microbial identification and metabolic profiling

Freshly grown (24-48 hr) colonies, from TSA or MRS agar, were used for microbial identification on GenIII microplates with the BIOLOG semi-automated system (Biolog, Hayward, CA) following the manufacturer’s instructions. Bacteria were added to the appropriate inoculating fluid and transmittance (T) was measured and adjusted to 90-98% T. For *Lactobacillus* species, Inoculating Fluid C (IF-C) was used. For *Bacillus* and other species, either Inoculating Fluid A or B (IF-A or IF-B) was used. Conditions are listed in **Table S1.** Suspended cells were dispensed with an automatic multichannel pipettor into the GenIII 96-well microplate (100 μl per well). The GenIII microplates were incubated at 33 °C for 16-48 hours and read using the MicroLog™ plate reader and associated software (Biolog, Hayward, CA) once the positive control well A10 turned purple (typically at 20-24 hr of incubation). Positive growth responses are indicated by a color change based on redox dye chemistry. Identification is made by the GENIII MicroStation™ software, which compares the phenotypic fingerprint with a fingerprint database of known bacteria [21]. Similarity (SIM) scores are assigned reflecting how well the isolate matches the pattern in the database, and an identification is given if the SIM score is >0.6. Bacterial identifications and SIM scores were recorded, and the plate image was saved for later analysis. The GenIII Microplate reference layout for each microorganism was also saved (see example in **Figure 2D**). The results from all Biolog plates were transcribed into a single summary table, using a “P” for positive reaction wells (purple), representing growth and utilization of a carbon source. Wells scored by the software as borderline (light color half-moon in Figure 2D, could be positive or negative) [21] were recorded as “h” for half in the results tables.

For yeast, colonies grown for 48-72 hour on SDA were inoculated into 10 ml sterile water, and the cell suspension was adjusted to 50% transmittance, then pipetted into the wells of a YT microplate.

YT plates were incubated at 26 °C for 24-72 hours, and analyzed at 24, 48, and 72 hours using the MicroLog™ plate reader until an identification was made.

### PCR amplification and sequencing of 16S rRNA gene

Polymerase Chain Reaction (PCR) was performed using DNA obtained directly from bacterial colonies. To amplify the near full-length 16S rRNA gene, the primers 27F (5’-AGAGTTTGATCCTGGCTCAG-3’) and 1492R (5’-GGTTACCTTGTTACGACTT-3’) were used. On ice, a Master mix was prepared, containing water, buffer, MgSO4, dNTPs, and primers following the manufacturer’s instructions for HotStart KOD Polymerase (Millipore Sigma, St. Louis, MO). Aliquots of 50 μl of the Master mix were added to labeled 8-strip PCR tubes, and a sterile 100 μl pipette tip was used to pick a pinpoint amount of a bacterial colony and transfer the cells directly into the appropriate PCR tube. Amplification reactions were run on a Bio-rad Thermocycler (Bio-rad, Hercules, CA) using the following cycling conditions: denaturation at 95 °C for 2 minutes, 35 cycles of: [95 °C denaturation for 20 seconds, 48 °C annealing for 20 seconds, 70 °C extension for 35 seconds], followed by a final extension step at 70 °C for 3 minutes. After visualizing the 1450-bp amplicons from each PCR reaction using gel electrophoresis, the PCR amplicons were purified using the QIAquick PCR Purification Kit (Qiagen, Valencia, CA). The DNA purified from the PCR reactions was quantified using a Take3 micro-volume plate with the Gen5 Microplate Reader (Biotek Instruments, Winooski, VT). DNA samples were prepared with forward or reverse primer and PCR products were sequenced by Sanger sequencing at the Yale University DNA Analysis Facility on Science Hill, using an Applied Biosystems Genetic Analyzer (New Haven, CT). The .abi files were downloaded and DNA chromatograms were viewed and trimmed using Geneious bioinformatics software (http://www.geneious.com/). Resulting forward and reverse sequences were searched against sequences in the Genbank non-redundant (nr) nucleotide database using Standard Nucleotide BLAST (blastn), and the top-scoring four hits were recorded for each organism.

### Matrix-assisted laser desorption ionization time-of-flight (MALDI-TOF) mass spectrometry

Bacteria from frozen stock cultures were transferred to MRS or TSA plates and incubated at 30°C for 48 hours prior to identification. Proteins were extracted using either the on-target method or by using an ethanol-formic acid protocol described by Friewald & Sauer (2009). Cells from isolated colonies were directly smeared onto a disposable FlexiMass™ DS target plate using a sterile toothpick. One μl of 25% formic acid was added to the spot and allowed to air dry followed by the addition of 1 μl of the α-Cyano-4-hydroxycinnamic acid (CHCA) matrix solution. The CHCA matrix solution contained 50 mg of CHCA dissolved in a 33/33/33 mixture of acetonitrile/ethanol/dH2O containing a final concentration of 3% trifluoroacetic acid. When the on-target method yielded spectra with poor resolution, proteins were extracted prior to spotting using ethanol and formic acid (Friewald & Sauer 2009). Cells from colonies were dissolved in 300 μl of dH_2_O and inactivated by adding 900 μl of room temperature absolute ethanol. The cell suspension was centrifuged twice at 10,000 x g for 2 minutes to remove the supernatant. The pellet was air dried at room temperature for 1 minute and dissolved in 10 μl 70% formic acid. Ten μl acetonitrile was added to the formic acid-cells mixture followed by centrifugation at 10,000 x g for 2 minutes at room temperature. The resulting supernatant containing extracted proteins was transferred to a separate tube. One μl of the supernatant was spotted onto the target plate and overlaid with 1 μl of the matrix.

MALDI-TOF MS was performed on the AXIMA Confidence iD^Plus^ MALDI-TOF Mass Spectrometer (Shimadzu) using Launchpad software version 2.9.1 and the VITEK^®^ MS Plus Spectral Archive and Microbial Identification System ™ (SARAMIS) database, V4.12. Samples were analyzed in the positive linear mode with a laser frequency of 50 Hz and within a mass range of 2000-20,000 Da.

The acceleration voltage was 20 kV and extraction delay time 200 ns. Spectra were generated from 500 laser shots and each target plate was calibrated before samples were analyzed using *Escherichia coli* DH5α. Samples were run in duplicate and spectra acquired by Launchpad were processed by SARAMIS. Each spectrum was assigned a confidence level based on a comparison to SuperSpectra in the SARAMIS database. SARAMIS does not assign a taxonomic name if the confidence levels are below 75%.

### Probiotic product label analysis and literature searches

To estimate the percentage of products listing specific strains on the label, products were evaluated from four marketplaces: a major online retailer, two drugstore chains, and a retail superstore with brick-and-mortar store locations in Shelton, CT. A search for products from the online retailer was conducted between Jan-April 2017 using the keyword “probiotic” and the name of the following probiotic microbes: *Bacillus coagulans, Bacillus subtilis, Lactobacillus acidophilus, Lactobacillus plantarum, Lactobacillus rhamnosus, Bifidobacterium,* and *Saccharomyces boulardii*. At least 20 products for each organism were checked, and if the label image contained a specific strain name or number, this was recorded. For brick-and-mortar stores (visited in April 2018), we counted unique products on the shelves in the probiotic section, examined the labels, and recorded the number of products that listed at least one strain ID on the label. In stores in which a high volume of store-brand generics was present, the products were only counted individually if the listed organisms were different from another store-brand product. Note that products found on the shelves at multiple stores were tallied each time in the count for that store; meaning that the list for each store often contained a high degree of overlap (particularly of the most popular name-brand probiotics).

To assess the representation of each type of probiotic bacteria and yeast in the medical literature, we performed literature searches of the Pubmed.gov database (https://www.ncbi.nlm.nih.gov/pubmed/) in March 2018. Keywords used were the genus and species name of each probiotic (for *B. coagulans,* we also included the former name, *Lactobacillus sporogenes*), AND “human.” The filter “Clinical Trial” was selected to count the number of published clinical trials.

## Results

### Isolation and culturing of probiotic microbes

Pure cultures of bacteria and yeast were isolated from 16 different probiotic supplements, six probiotic beverages, and four environmental sources (Chaas fermented beverage, fruit fly gut, kale, and a leaf). Though most probiotic microorganisms are facultative anaerobes, their aerobic growth was comparable to growth with CO_2_ Gaspaks, and further experiments were conducted under aerobic conditions. *Bifidobacterium* spp. were excluded from this investigation due to their strict anaerobic and fastidious growth. The main bacterial species cultured were *B. coagulans, B. subtilis, L. plantarum,* and *L. rhamnosus,* with typical colony morphology on TSA or MRS agar under aerobic conditions shown in **Figure 1.** While the *Lactobacillus* species have very similar colony appearance (off-white to white, circular, creamy), the two *Bacillus* species are quite distinct, with *B. coagulans* colonies showing irregular edges, translucent tan color, and less robust growth. The colonies of *B. subtilis* have more rapid, spreading growth, and are opaque off-white in color, raised, and wrinkled. Gram staining confirmed that all bacteria were Gram-positive rods. As expected, spore staining revealed that only the *Bacillus* species formed spores (not shown). *B. coagulans* formed visibly stained spores only when the bacteria were grown on TSA, not on MRS. In one single-strain probiotic labeled as “*Bacillus coagulans,*” both *B. coagulans* and *B. subtilis* were cultured repeatedly from the capsule, indicating contamination within the original product.

**Figure 1.**
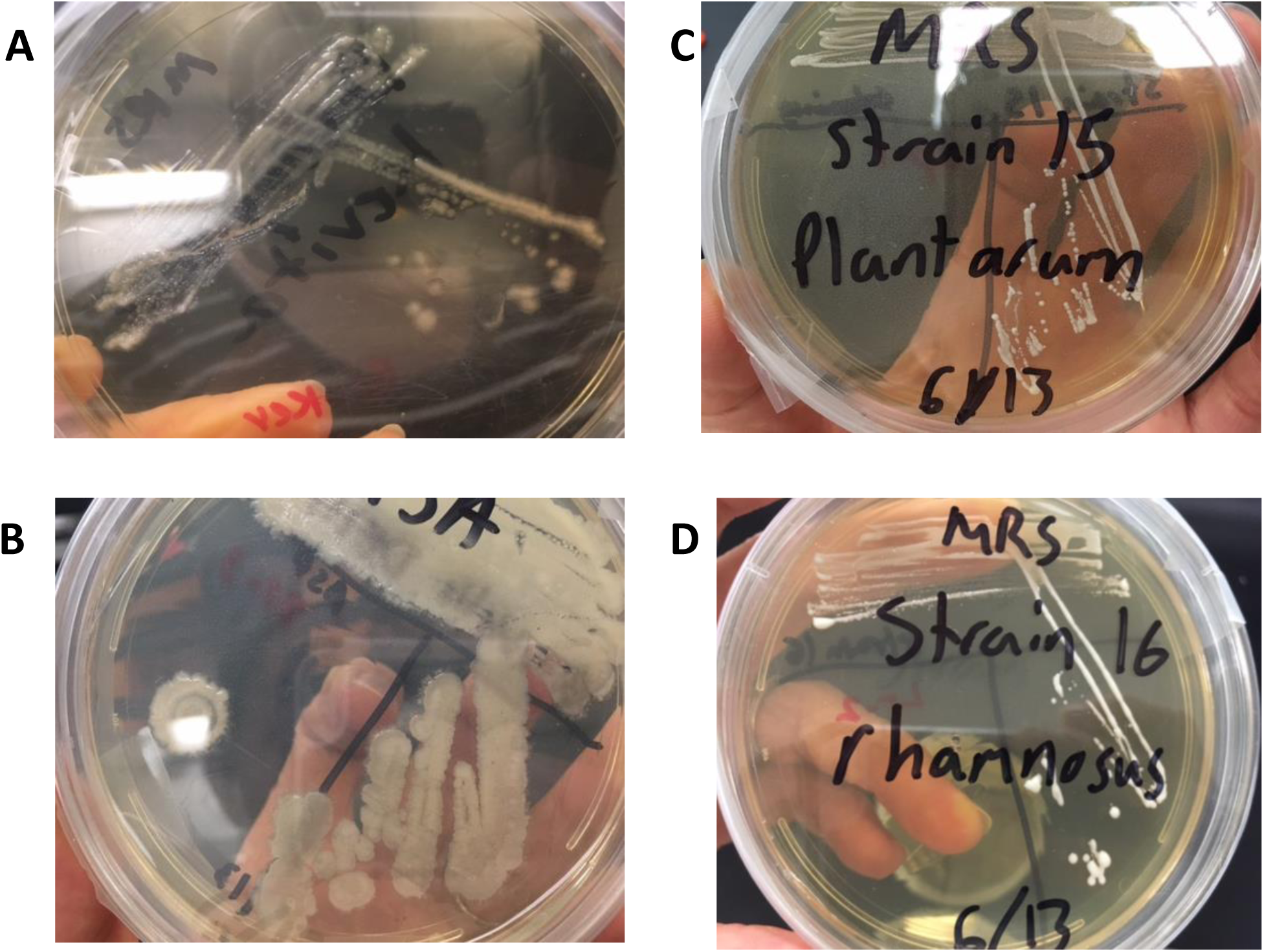
Typical colony morphology of probiotic bacterial colonies grown on agar media. A: *Bacillus coagulans* on MRS agar, B: *Bacillus subtilis* on TSA, C: *Lactobacillus plantarum* on MRS agar, D: *Lactobacillus rhamnosus*. on MRS agar.

### Identification of isolated microbes

**Table 1** summarizes the number of identifications, obtained using each of the three methods, that matched the listed species on the label.

**Table 1.** **Number of correct bacterial species identifications by each method**

| Name | # of Products tested | Correct ID with 16S sequencing | Correct ID with MALDI | Correct ID with 278 BIOLOG 279 |
| --- | --- | --- | --- | --- |
| *Bacillus coagulans* | 4 | 3 | 2 | 0 280 |
| *Bacillus subtilis* | 3 | 2 | 3 | 3 281 |
| *Lactobacillus plantarum* | 5 | 4 | 3 | 5 |
| *Lactobacillus rhamnosus* | 4 | 3 | 4 | 3 282 |
| Other *Lactobacillus* spp. | 2 | 2 | 0 | 0 |
| Total | 18 | 14 (78%) | 12 (67%) | 11 (61%) 283 |

The species identifications obtained by each method, for each isolate, are listed in **Table 2.** Each isolate was assigned a Code number to de-identify the product brand. Table 2 includes a total of 26 isolated organisms. For the *S. boulardii* probiotic yeast, the only method used was Biolog identification, and two out of three products listing *S. boulardii* were correctly identified with the Biolog system. This total exceeds the 18 total products listed in Table 1, because it includes microbes for which the identity was unknown prior to analysis (some probiotic yeasts, and environmental isolates from the Chaas fermented beverage, fruit fly gut, and leaves). Seven of the products listed the specific strain identification (Strain ID) on the label: *B. coagulans* GBI-30 6086 (patented as GanedenBc^30®^), *B. subtilis* DE111, *L. plantarum* 299V, *L. rhamnosus* GG, and *L. rhamnosus* LCR35 **(Table 2)**.

**Table 2.** **Species identifications of probiotic microbes using three methods**

| Source | Code | Label Identification | 16S sequence BLAST Result | MALDI Result | Biolog Result |
| --- | --- | --- | --- | --- | --- |
| *Bacillus coagulans* |  |  |  |  |  |
| Probiotic Beverage | 5 | *Bacillus coagulans*<br>GBi-30 6086 | *Bacillus coagulans* | *Bacillus coagulans* | No ID |
| Probiotic Beverage | 6 | *Bacillus coagulans*<br>GBi-30 6086 | *Bacillus coagulans* | *Bacillus subtilis* | *Sporolactobacillus*<br>*kofuensis* |
| Single-strain Probiotic | 8 | *Bacillus coagulans* | contamination:<br>*Bacillus subtilis* ** | *Bacillus coagulans* | *Lactobacillus*<br>*paracasei* |
| Single-strain Probiotic | 14 | *Bacillus coagulans*<br>GBi-30 6086 | *Bacillus coagulans* | *Lactobacillus*<br>*pentosus/plantarum* | *Brochothrix*<br>*campestris* |
| ***Bacillus subtilis*** |  |  |  |  |  |
| Single-strain Probiotic | 9 | *Bacillus subtilis* | *Bacillus* sp. strain<br>BCBT29 | *Bacillus subtilis* | *Bacillus*<br>*atrophaeus/subtilis* |
| Multi-strain Probiotic | 13 | *Bacillus subtilis* | *Bacillus subtilis* | *Bacillus subtilis* | *Bacillus subtilis* |
| Multi-strain Probiotic | 15 | *Bacillus subtilis*<br>DE111 | *Bacillus subtilis* | *Bacillus subtilis* | *Bacillus subtilis* |
| ENV: leaf surface | 20 | N/A (environmental) | N/A | N/A | *Bacillus*<br>*atrophaeus/subtilis* |
| ***Lactobacillus***<br>***plantarum*** |  |  |  |  |  |
| Single-strain Probiotic | 2 | *Lactobacillus*<br>*pentosus/plantarum* | *Lactobacillus*<br>*plantarum* | *Lactobacillus*<br>*pentosus/plantarum* | *Lactobacillus*<br>*plantarum* |
| Multi-strain Probiotic | 11 | *Lactobacillus*<br>*plantarum* complex | *Lactobacillus*<br>*plantarum* | No ID | *Lactobacillus*<br>*plantarum* |
| Multi-strain Probiotic | 12 | *Lactobacillus*<br>*plantarum* | *Lactobacillus*<br>*plantarum* | *Lactobacillus*<br>*plantarum* | *Lactobacillus*<br>*plantarum* |
| Probiotic Beverage | 16 | *Lactobacillus*<br>*plantarum* 299V | *Lactobacillus*<br>*plantarum* | *Lactobacillus*<br>*pentosus/plantarum* | *Lactobacillus*<br>*plantarum* |
| ENV: Fruit fly gut | 19 | N/A (environmental) | *Lactobacillus*<br>*plantarum* | N/A | *Lactobacillus*<br>*plantarum* |
| ***Lactobacillus***<br>***rhamnosus*** |  |  |  |  |  |
| Single-strain Probiotic | 1 | *Lactobacillus*<br>*rhamnosus* GG | *Lactobacillus*<br>*rhamnosus* | *Lactobacillus*<br>*rhamnosus* | *Weissella viridescens* |
| Multi-strain Probiotic | 7 | *Lactobacillus*<br>*rhamnosus* | *contamination: *S.*<br>*epidermidis* | *Lactobacillus*<br>*rhamnosus* | *Lactobacillus*<br>*rhamnosus* |
| Single-strain Probiotic | 10 | *Lactobacillus*<br>*rhamnosus* LCR35 | *Lactobacillus*<br>*rhamnosus* | *Lactobacillus*<br>*rhamnosus* | *Lactobacillus*<br>*rhamnosus* |
| Multi-strain Probiotic | 17 | *Lactobacillus*<br>*rhamnosus* | *Lactobacillus*<br>*rhamnosus* | *Lactobacillus*<br>*rhamnosus* | *Lactobacillus*<br>*rhamnosus* |
| ENV: fermented milk<br>beverage "Chaas" | 18 | N/A (environmental) | *Lactobacillus casei* | *Lactobacillus* sp. | *Lactobacillus*<br>*rhamnosus* |
| **Other *Lactobacillus***<br>***spp.*** |  |  |  |  |  |
| Single-strain Probiotic | 3 | *Lactobacillus gasseri* | *Lactobacillus*<br>*gasseri* | No ID | *Streptococcus oralis*<br>or *Paenibacillus*<br>*sanguinis* |
| Single-strain Probiotic | 4 | *Lactobacillus acidophilus* | *Lactobacillus acidophilus* | No ID | No ID |
| **YEAST** |  |  |  |  |  |
| Kombucha 1 | 21 | *Saccharomyces boulardii* | N/A | N/A | *Saccharomyces cerevisiae a/Tor.pretorein* |
| Kombucha 2 | 22 | *Saccharomyces boulardii* | N/A | N/A | *Pichia chmeri A* |
| Kombucha 3 | 23 | N/A | N/A | N/A | *Hanseniospora guillermundii/uvarum* |
| Probiotic rice wine | 24 | *Saccharomyces cerevisiae* | N/A | N/A | *Saccharomyces boulardii* |
| Multi-strain probiotic | 25 | *Saccharomyces boulardii* | N/A | N/A | *Saccharomyces boulardii* |
| ENV: Kale leaf | 26 | N/A (environmental) | N/A | N/A | *Sporidiobolus pararoseus A* |

### Identification by 16S rRNA gene sequencing

Molecular identification by sequencing the 16S rRNA gene is currently a routine method of confirming species identity. The universal primers 27F and 1492R were used to amplify the near full length 16S gene, using direct colony PCR. Since this gene sequence is generally not variable enough to distinguish between strains of the same species, we used 16S rRNA sequencing primarily as a quality control check to ensure that we had indeed isolated the correct species from each product. For those products listing the Strain ID on the label, the top-scoring BLAST hit of the 16S sequence usually did not match the exact strain name (with the exception of *L. rhamnosus* strain “GG.”), and the % sequence identity of the five top-scoring BLAST hits were typically identical **(Table S2)**. In most cases, the nucleotide BLAST search of the forward and the reverse sequences yielded different strain names of the expected species **(Table S2)**. Several of the incorrect IDs (strain codes 7 and 8) may be attributed to contamination during the procedure (Table 2).

### MALDI-TOF identification

Using Matrix Assisted Laser Desorption Ionization Time of Flight Technology Mass Spectroscopy (MALDI-TOF), the mass fingerprints of 15 bacteria isolated from commercial probiotics or environmental sources were obtained and identified at the genus and species level. For 12 out of 18 isolated strains the MALDI identification matched that of the probiotic label **(Tables 1+2)**. The “Chaas” isolate was a homemade fermentation so the bacterial identity was not labelled; MALDI-TOF MS identified this strain only on the genus level (*(Lactobacillus sp*.). All *Bacillus subtilis* isolates were correctly identified by MALDI-TOF MS. *Bacillus coagulans* strains were not as consistently identified by this technique. The probiotic drink isolate (strain 14) was identified as *Lactobacillus pentous/plantarum*, while the probiotic label indicated the strain was *Bacillus coagulans. Bacillus coagulans* (strain 6) was misidentified by MALDI as *B. subtilis*. Several *Lactobacillus* strains were unable to be identified: *L. acidophilus, L. gasseri,* and one *L. plantarum* (strain 11.) However, other *L. plantarum* isolates were successfully identified so this species is represented in the SARAMIS database. If a given peptide mass spectrum matched the database with a confidence score <75%, no identification in SARAMIS could be reported; confidence interval scores are provided in **Table S3**.

### Biolog Identification & Phenotypic Profiling

Of the 18 probiotic bacterial isolates, 11 were correctly identified using the Biolog Microbial ID system **(Table 1, Table 2)**. For 17 of these strains, the expected identity of the bacteria was given on the product label, however the strain isolated from the “Chaas” fermented milk beverage was not known in advance. Among the *Lactobacillus* isolates, *Lactobacillus plantarum* strains were the most amenable to identification with the Biolog system. All of the *L. plantarum* strains isolated from probiotic products were correctly identified, with SIM scores >0.6 **(Table S1)**. One *L. rhamnosus* strain yielded an incorrect identification, erroneously reading as *Weissella viridescens*. Neither *L. acidophilus* nor *L. gasseri* were identified correctly with the Biolog assay. For the *Bacillus* strains isolated from probiotics, *Bacillus subtilis* was more readily identified using the Biolog system. Three *B. subtilis* isolates from different products were accurately identified, while none of the three *B. coagulans* isolates came up as *B. coagulans*. Factors such as poor growth, use of non-optimal inoculating fluid or medium, or inappropriate incubation conditions may have had adverse effects on the phenotypic expression pattern. The time of incubation of the GenIII microplate is critical to a correct identification, as is the growth medium. While the Biolog manufacturer recommends BUG-B agar, we observed better growth of *B. coagulans* on MRS medium, so MRS was used.

**Figure 2** summarizes the results from the Biolog phenotypic profiling for each strain, with carbon source utilization patterns shown in Figure 2A and tolerance to environmental stressors (acidity, salt, and various compounds) in Figure 2B. The reference pattern stored in the Biolog GenIII database is displayed in the top row for each species. The darker color “P” wells indicate a strong positive, while the lighter “h” wells indicate borderline results (Figure 2D); the preferred carbon sources are typically used up more rapidly and completely, yielding a dark purple well, while less preferred substrates may be used more slowly and incompletely[21]. Results of antibiotic susceptibility testing for 12 antibiotics using the disc diffusion (Kirby-Bauer) method are also included in Figure 2. Strain-specific differences were observed for all of the species, in both carbon source utilization and environmental stress tolerance.

**Figure 2.**
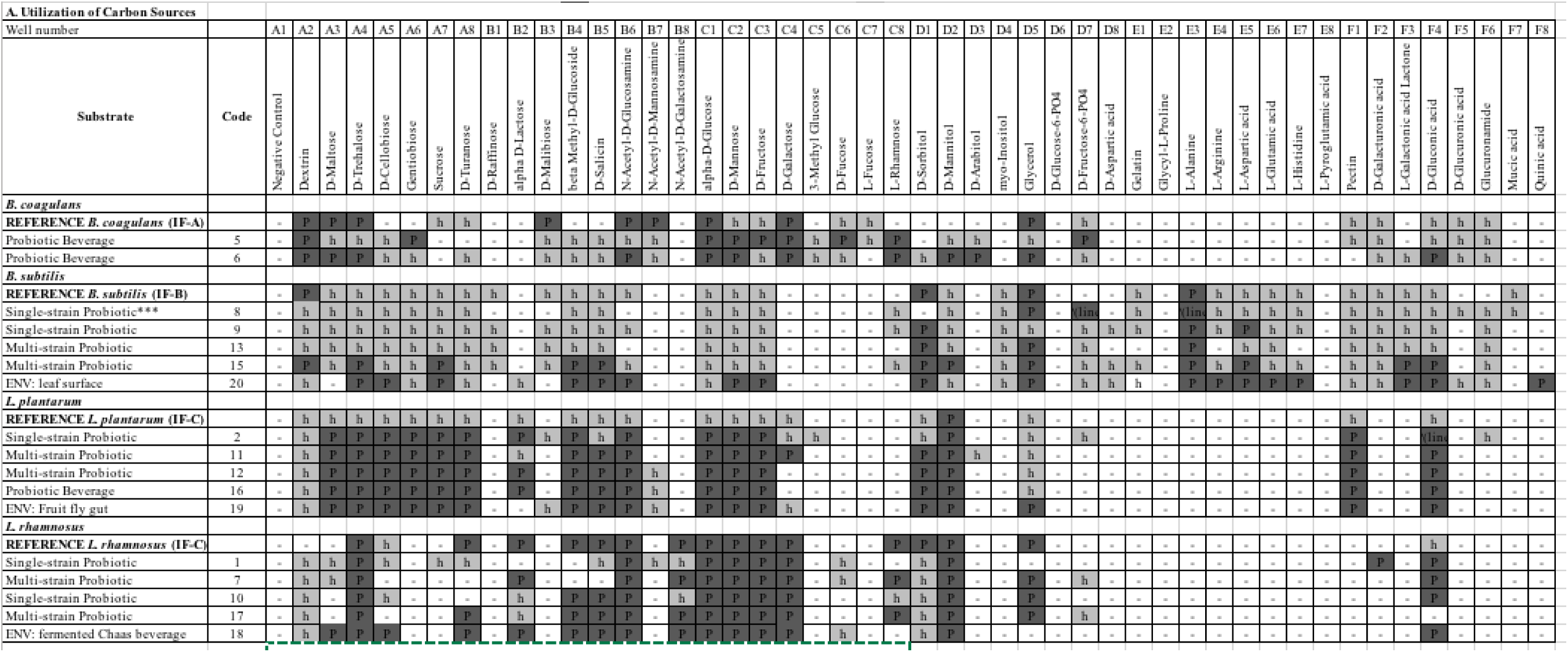
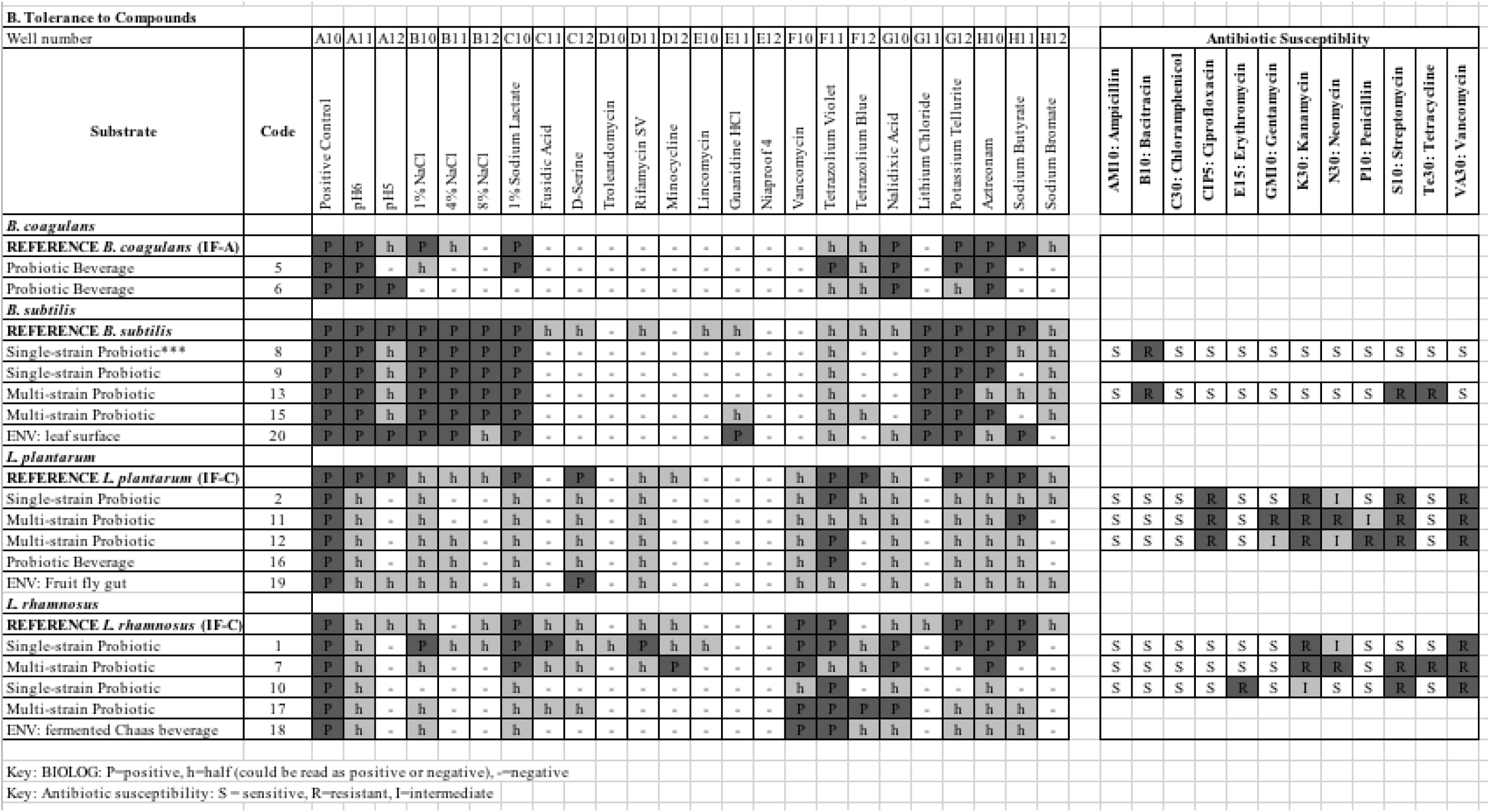
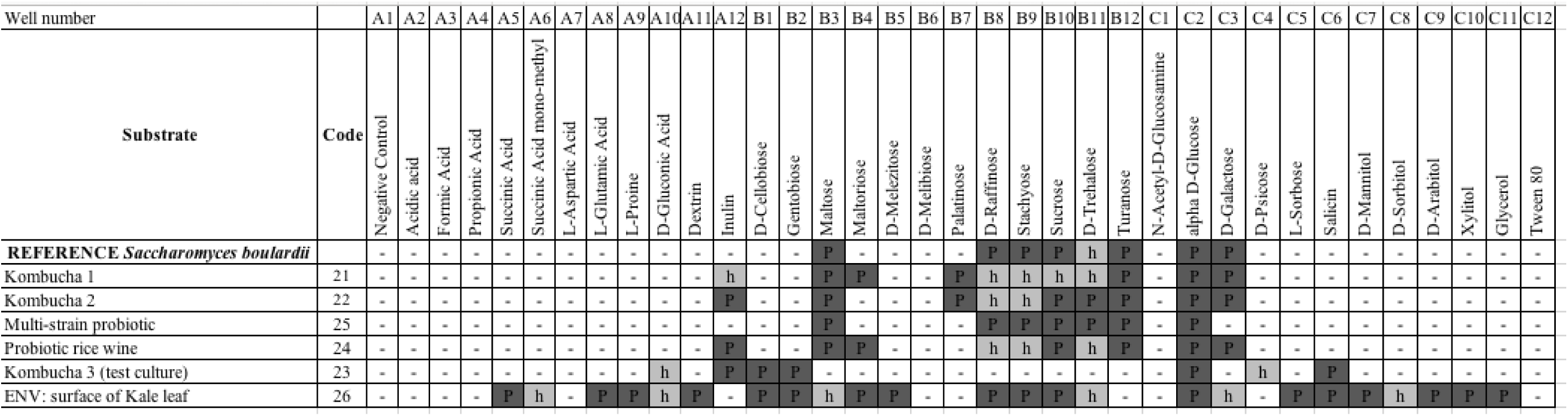
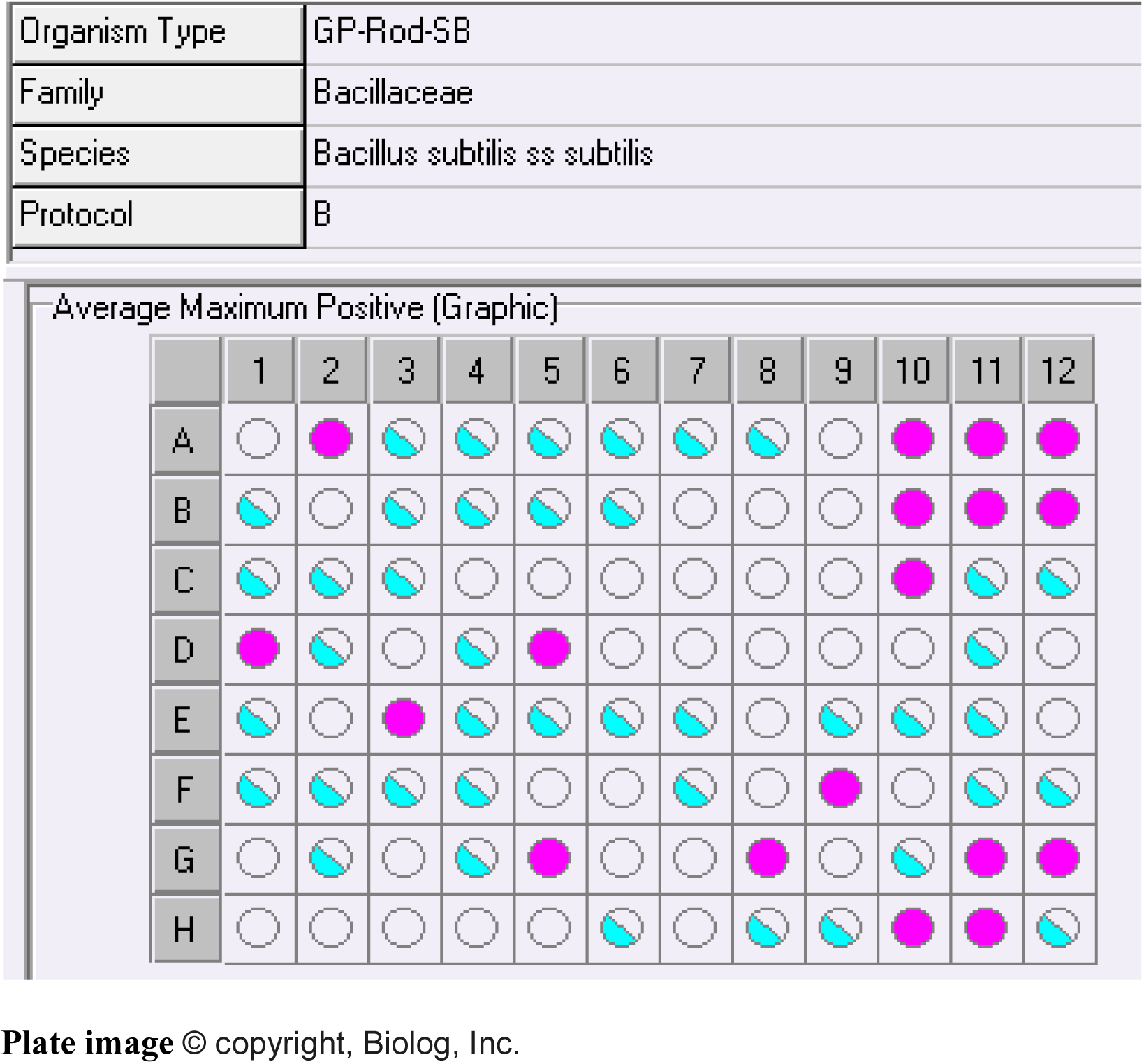
Phenotypic profiling of probiotic bacteria and yeast with Biolog system. P=positive, h=half (borderline positive/negative). **A**: Utilization of carbon sources. **B:** Tolerance to compounds, including antibiotic susceptibility. **C:** Carbon source utilization by yeasts isolated from probiotic drinks. Note that wells on the yeast (YT) plate in C contain different carbon sources than the plate for bacteria (GenIII). **D:** Example Biolog plate diagram showing the reference pattern for *Bacillus subtilis,* pink=positive; half-moon=borderline.

The Biolog system is also capable of identifying fungi such as yeast, using specific YT yeast plates and the YT database. After optimizing the time and temperature (72 hours at 26 °C), we identified yeast strains isolated from four probiotic drinks, one probiotic supplement and the surface of a kale leaf with Biolog YT **(Table 2, Figure 2C)**. Three of these (strain codes 21,22, 25) were labelled as containing *Saccharomyces boulardii,* and the correct identification was obtained for two out of three.

The metabolic utilization patterns of 35 carbon sources were compared for these probiotic yeasts and can be viewed in **Figure 2C.** The sugars glucose (Well D6/D7), turanose (Well A8), and maltose (A3) had clear positive growth in the all *S. boulardii* isolates. Inulin (Well A12), a common prebiotic fiber, was utilized by several strains, although the reference pattern for *S. boulardii* used in the Biolog database is negative for inulin (Figure 2C).

### Evaluation of product labels and quantification of published clinical studies

One objective of this project was to estimate the percentage of probiotic products currently on the market that list the specific strain ID of the bacterial or yeast species on the ingredient label.

Products available from a major online retailer, at two drugstore chains, and at a retail superstore were evaluated by reading the ingredient label and recording whether or not each product listed an alphanumeric strain ID (often the patented name of the strain). **Table 3** summarizes these product counts from four retail sources. With this approximation we saw that an average of 49% (ranging from 34-69%) of products contained specific strain information on the label.

**Table 3.** **Strain identifications listed on probiotic product labels at retail stores**

| Store | # products checked | # products with Strain ID | % with Strain ID | % without Strain ID |
| --- | --- | --- | --- | --- |
| Major online retailer | 121 | 41 | 34% | 66% |
| Drugstore 1 | 28 | 13 | 46% | 54% |
| Drugstore 2 | 21 | 10 | 48% | 52% |
| Retail superstore | 26 | 18 | 69% | 31% |

To examine the number of clinical studies performed on these probiotic strains, providing context for the amount of evidence supporting their health claims, we searched the biomedical literature database Pubmed.gov, for clinical trial publications on each species. **Figure 3** shows the total counts of published clinical trials for eight common probiotic microbes. The well-established probiotic species, *Lactobacillus acidophilus*, *Bifidobacterium longum*, and *Bifidobacterium infantis* (a substrain of *B. longum*) were included in this literature search despite not being characterized in the current study. As shown in **Figure 3,** *L. acidophilus* and *L. rhamnosus* have each been studied in over 300 clinical trials to date. In contrast, there are fewer clinical trials published on *Bacillus* probiotics (49 found for *B. coagulans* and 47 for *B. subtilis*). Evaluation of the content or quality of these studies was not performed.

**Figure 3.**
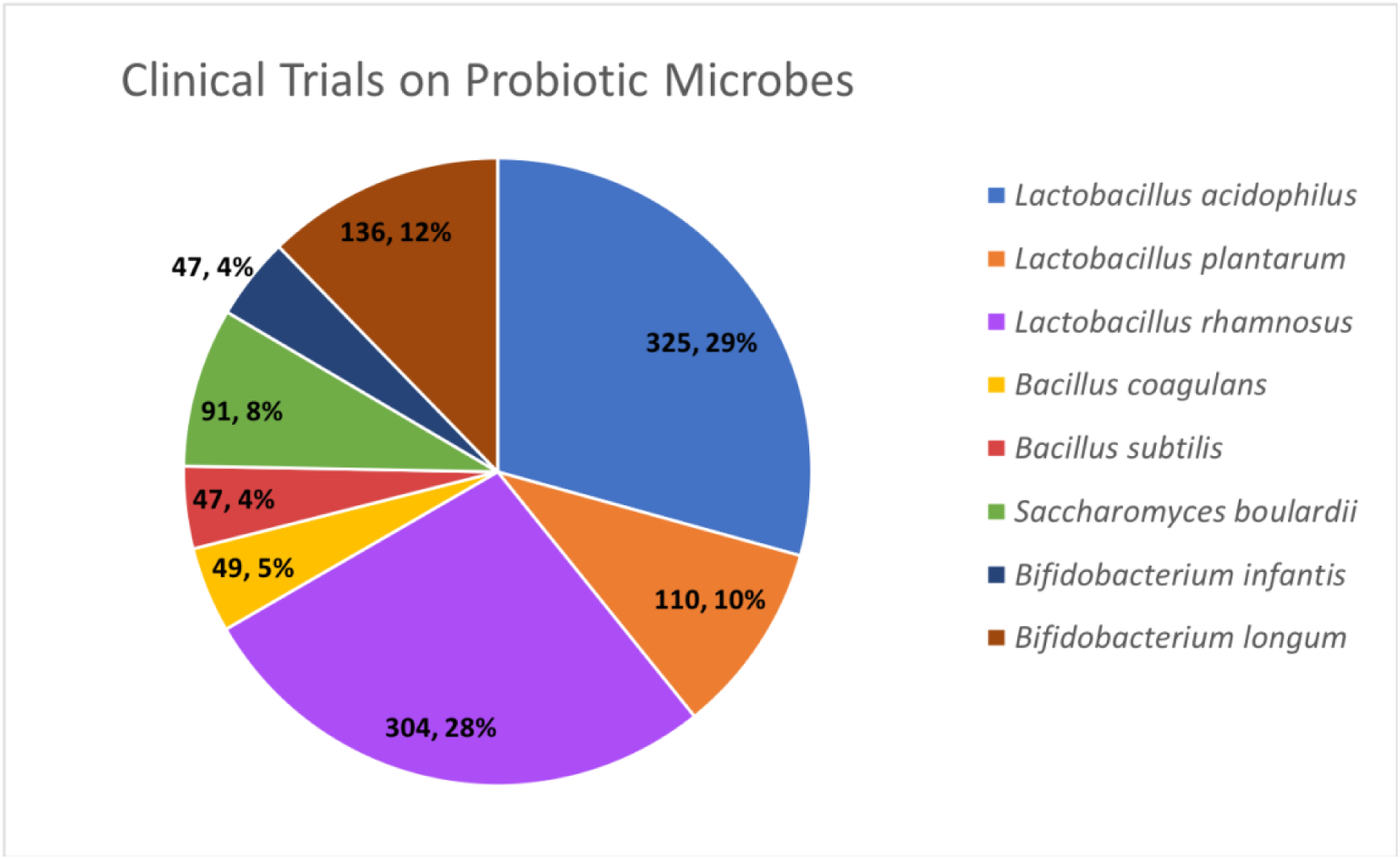
Number of published clinical trials on eight major probiotic species, cataloged in the Pubmed.gov database as of March 2018.

## Discussion

We isolated and cultured microbes from commercially available probiotic supplements and beverages and used three approaches to identify the bacteria from these products. We then looked for evidence of strain-level diversity between the isolates using phenotypic profiling. Sequencing of the 16S gene and MALDI-TOF mass spectrometry both yielded accurate identifications for a higher percentage of bacteria than the Biolog assay. Sequencing of the 16S rRNA gene has become a standard molecular identification technique since its introduction in the 1980s [22], but is generally not sufficient for strain typing. MALDI-TOF mass spectrometry is a newer method for rapid identification of bacteria, however it requires that the peptide mass spectra already exist in the database [23]. When comparing 16S sequencing, Biolog identification, and other methods for identifying *Lactobacillus* species, only MALDI mass spectrometry was noted for the ability to identify species at the subspecies, or strain level [24]. A study on 148 strains of *Lactobacillus* species isolated from food found that the MALDI technique generated accurate species identifications more often than 16S PCR (93% accuracy vs. 77% for PCR), and the authors suggest that MALDI should be used in combination with genotypic methods for improved reliability [25]. Sato and colleagues (2017) used MALDI-TOF and repetitive sequence based PCR (rep-PCR) for rapid strain typing of strains of *B. coagulans* [26]. This group found a strong correlation between these two methods to successfully distinguish between closely related strains, and reported that carbohydrate utilization patterns correlated well with the MALDI and rep-PCR results for some phylogenetic clusters [26]. We observed agreement in the results from the three methods, with minor exceptions. For strain 1, MALDI and 16S results were in agreement, correctly identifying *L. rhamnosus* **(Table 2)**, but the Biolog identification of this isolate was *Weissella viridescens,* a lactic acid bacterium (formerly classified as a *Lactobacillus*), often found in fermented foods [27]. Its pattern of carbon source usage is quite similar to the other *L. rhamnosus* strains. For the “Chaas” isolate, 16S sequencing identified this as *L. casei,* while Biolog suggested *L. rhamnosus*. These species are closely related and belong to the *L. casei* group along with *L. paracasei*.

Several other molecular approaches have been used for distinguishing bacteria at the strain level, such as multi-locus sequence typing (MLST), pulsed-field gel electrophoresis (PFGE), or amplified fragment length polymorphism (AFLP). MLST used to be a “gold standard” for differentiating between strains, however this is changing as costs of whole-genome sequencing (WGS) have decreased [28]. PFGE and AFLP have been used to differentiate among probiotic strains of *L. rhamnosus* [29] and *L.plantarum* isolated from various sources [30]. Ceapa and colleagues (2015) identified genotypic clusters of *L. rhamnosus* with AFLP that correlated with functional metabolic clusters determined by Biolog profiling [31]. Due to decreasing costs and improved efficiency, WGS is likely to be the future of microbial strain typing [32]. Phylogenomics approaches have recently been proposed to improve the classification of the diverse *Lactobacillus* genus [33,34], and metagenomic strategies with bioinformatics analyses are in being developed to characterize microbial diversity at the subspecies level in human microbiome communities [35] [36].

Several studies have used comparative genomics to investigate the genetic basis of the probiotic properties and predicted metabolic capabilities of these organisms. Khatri and colleagues compared genomes of *Bacillus coagulans* to *Bacillus subtilis,* and found considerable genetic heterogeneity in *B. coagulans* strains [37], which has been recapitulated in carbohydrate utilization assays [26]. Analyzing the genome of the commercialized *B. coagulans* strain HM-08 uncovered a xylose utilization gene cluster, lending insight to future biotechnological applications for lactic acid production by this strain [38]. Genome analysis has revolutionized the classification and characterization of lactic acid bacteria, and functional genomics investigation has led to the discovery of novel processes of communication with host cells, that may serve as models for understanding other host-microbe dynamics [39]. Comparative genomics of *Bifidobacterium* species has unveiled substantial diversity carbohydrate metabolism [40] and helped identify the cell surface proteins and exopolysaccharides used to colonize the host intestine [41]. The wide range of polysaccharides used by *B. longum* species has been proposed to aid in their success as early colonizers of the infant gut [42]. By comparing the genome of *Lactobacillus rhamnosus* GG to a less-adherent starter culture strain, a unique genomic island encoding secreted pilins that bind to human intestinal mucus was discovered [43]. These mucus-binding pilins were shown to specifically outcompete vancomycin-resistant *Enterococcus faecium* (VRE) due to homology in the binding site [44].

Despite its lower performance to correctly identify probiotic bacteria, the Biolog 96-well plate assay is a valuable tool for rapidly collecting phenotypic data on 71 carbon sources and 23 environmental stressors in one multiplex assay, and illustrates a critical link between prebiotics and probiotics. By comparing the carbon source utilization patterns of the *Lactobacillus* and *Bacillus* species, we can gain insight on shared and unique metabolic properties of these probiotic strains. The most striking difference between the two *Lactobacillus* species compared to the *Bacillus* species, is their overwhelming preference for sugars. *Lactobacillus plantarum* strains showed a very consistent pattern of sugar utilization, with 14 sugar wells having a positive reaction for all five of the strains tested. These results correlate with comparative functional genomics and metabolic profiling studies on *L. plantarum* [30][45]. *L. plantarum* probiotic strains, but not *L. rhamnosus,* utilized the complex polysaccharide, pectin (Well F1), found in the skins of many fruits. This may be related to its prevalence in plant-associated habitats. Pectinolytic enzymes have been characterized in *L. plantarum* [46] and pectin affects the probiotic phenotype of this species *in vitro* [47]. Likewise, *L. plantarum* grew on gentobiose (Well A6), a rare disaccharide found in the gentian family of plants. Only *L. rhamsosus* utilized the sugar rhamnose (Well C8). The antibiotic susceptibility profiles of *L. rhamnosus* vs. *L. plantarum* are slightly different **(Figure 2)**. Like most lactobacilli, they are naturally resistant to vancomycin (Well F10) due to absence of D-ala in the peptide crossbridge of their cell walls [48], but *L. plantarum* strains also displayed resistance to ciprofloxacin. The three *L. rhamnosus* strains tested were susceptible to penicillin.

As a soil microbe subject to nutrient limitation in the environment, *B. subtilis* is known to be much more versatile in its metabolism [49], and this is reflected in its Biolog metabolic profiles. A variety of amino acids are utilized, such as L-alanine (Well E3) and L-aspartic acid (Well E5) **(Figure 2A)**. These wells turned purple (positive) more rapidly than some of the sugar wells. *B. subtilis* is the most salt-tolerant of the species investigated, growing at up to 8% NaCl (Well B12), but is more susceptible to antibiotics than the lactobacilli **(Figure 2B)**. *Bacillus coagulans,* formerly classified as *Lactobacillus sporogenes* and isolated from spoiled milk in 1915, inhabits ecological niches more similar to lactic acid bacteria than other *Bacillus* spp. [50]. Although the Biolog approach did not correctly identify the *B. coagulans* isolates, their pattern of carbon metabolism is primarily metabolizing sugars, similar to the *Lactobacillus* species, and with the ability to withstand pH 5 (Well A12, **Figure 2B)**. While *B. coagulans* is categorized as a “GRAS” ingredient, safety considerations are critical for consumption of *Bacillus subtilis* [15]. It is unclear what the source of *B. subtilis* lacking Strain IDs are, as the Biolog ID of strain 9 (from a single strain probiotic also containing plant extracts) resulted in the same identification, *Bacillus atrophaeus/subtilis,* as the ID for a *B. subtilis* environmental isolate, which we cultured from the surface of a leaf. The profile of this environmental *B. subtilis* was quite similar to the strains from probiotic supplements, with the exception of utilization of quinic acid (well F8), a compound found in plant sources. The spore-forming ability of *Bacillus* species makes them highly stable probiotics that can be easily added to food or gummy supplements, however *B. subtilis* especially merits increased regulation for safety and efficacy.

It is well known that strain-level differences occur in the probiotic properties of microorganisms [51]. From survival in the GI tract (by tolerance to acidic pH and bile salts), to adhesive capacity to intestinal cells, to competition with pathogens and production of bioactive compounds, the capacity and efficiency to perform these functions is often strain-dependent [52][53]. Here we show strain-level differences in several environmental stressors, such as salt tolerance **(Figure 2B,** wells B10-B12) among the *Lactobacillus* strains, and considerable variation in nutritional phenotypes, based on profiling of carbon source usage. There is evidence that these two phenomena are related: the food sources and molecular cues that microbes encounter in their environment affect their expression of proteins and compounds (or community-level behavior such as aggregation and biofilm formation) that convey the probiotic’s beneficial effect. Further experiments are needed to measure a correlation between the metabolic profiles and probiotic properties of these particular strains, similar to a study of two lactic acid bacteria from the commercial culture FloraMax^®^-B11 [54]. Several examples of the relationship between nutrient sources and bacterial probiotic phenotype include: increased resistance of *L. plantarum* to gastric juices when growth with pectin or inulin compared to glucose [47]; differences in cell surface hydrophobicity, cell surface protein and exopolysaccharide production of *L. rhamnosus* grown on fructose, mannose, or rhamnose [55]; and increased the adhesion of *Lactobacillus acidophilus* to mucin or intestinal cells in the presence of fructooligosaccharides (FOS), cellobiose, or polydextrose [56]. The prebiotic cellobiose was shown to change surface layer proteins and increase auto-aggregation in two *Lactobacillus* strains [57].

A thorough understanding of the nutritional preferences of the commensal bacteria for current and next-gen probiotics will be vital for their translation into effective products, and metabolic profiling can help inform the design of “synbiotic” foods (containing probiotics and prebiotics) [58][6] and “biofunctional” foods (in which microorganisms cause the desired biological or physiological effect) [51]. If end-products of microbial metabolism contribute to the health-promoting effect, it will be imperative that the target microbes have an ample supply of and access to the carbon sources and environmental signals that lead to synthesis of those end-products. For example, plant glucosides from fruit are metabolized by *L. acidophilus,* which then secrete aglycones that exert beneficial effects on the host [59]. Among the wide array of bioactive compounds produced by lactic acid bacteria are B vitamins, gamma-aminobutyric acid (GABA), bioactive peptides, bacteriocins, and other complex molecules such as exopolysaccharides [51]. Several other recent review articles summarize the health benefits provided by microorganisms in functional foods [60] and explore the idea of how dietary composition can reshape a healthy microbiome to restore functions lost through the Western diet and lifestyle [61][62][7]. In a randomized clinical trial evaluating dietary interventions for type 2 diabetes mellitus, a high fiber diet altered the composition of the gut microbiota, and greater diversity of carbohydrates was associated with improved clinical outcomes [63]. The authors noted strain-specific effects on which active SFCA-producing bacteria were positive responders to the fiber, such as certain strains of *Faecalibacterium prausnitzii* [63].

While traditional probiotic bacteria (lactobacilli and bifidobacteria) have been the subject of research for decades, less is known about the preferred nutritional requirements of other dominant members of the human gut microbiome. Using defined media, Tramontano and colleagues tested the carbon source utilization of gut commensals (including mucin, carbohydrates, and the inhibition of growth of some by amino acids or SCFAs [64]. ‘Culturomics’ studies combining MALDI-TOF identification with >200 culture conditions optimized for fastidious growth have led to identification of 341 species of bacteria cultured from stool samples and helped optimize culture conditions for these bacteria [65]. Better models to study the gut ecosystem are needed, to understand the context of the metabolic interactions between gut commensals and probiotics.

In conclusion, there is currently a disparity between the composition of marketed probiotics available to consumers and the science backing their claims **(Figure 3)**. In the food supplement industry, the specific strain of a species is not always designated on the label, particularly for the generic, store-brand, or less-established brands. This may be due in part to proprietary restrictions. We found that on average, roughly half of the probiotics examined had the specific strain listed on the label, which varied considerably by store **(Table 3)**. Regulatory guidelines differ widely across different countries [66]. Probiotic labeling at the strain level is critical to so that consumers and/or healthcare providers can more easily evaluate clinical studies of the probiotic’s effects for specific indications [13], even if these properties are shared among all members of a species [16]. This study of common commercially available probiotics highlights the importance of supporting health claims by correctly identifying the microbes in probiotics, and the importance of understanding the ecophysiological needs of a given microbe to enable its beneficial effect (e.g. competition, colonization, flocculation, biofilm, antibiotic production, adhesion, bioactive metabolite production). In the complex milieu of the digestive tract, metabolic profiling of individual microbes and microbial communities can help draw the link between prebiotics, probiotics, and overall digestive health. These insights could bring about strategies for optimizing health and wellness, grounded in nutrition that promotes synergy with our commensal microbiota.

## Acknowledgments

We thank Fairfield University biology students in the Fundamentals of Microbiology Lab (Fall 2016) for their contributions to the project, including isolating and characterizing probiotic bacterial strains. We are grateful for assistance from Eunsun Hong, Jenna Massaro, Samantha Porter and Philip Strang, with PCR and BIOLOG microbial identification, and to the Fairfield University Biology lab supervisors Christopher Hetherington and Lenka Biardi for their valuable assistance. We would like to thank Carol Mariani at the Yale University DNA Analysis Facility on Science Hill (New Haven, CT) for assistance with DNA chromatogram interpretation. We also thank Dr. Jillian Smith-Carpenter of the Department of Chemistry & Biochemistry at Fairfield University for guidance and training on the MALDI-TOF instrument, supported by the NSF grant CHE-1624744 awarded to Fairfield University.

